# Cytoprotective effects of combination of gypenosides and *Costus pictus* D.Don extract in BRIN-BD11 β-cells

**DOI:** 10.1101/2020.09.07.286435

**Authors:** Chinmai Patibandla, Mark James Campbell, Leigh Ann Bennett, Xinhua Shu, Steven Patterson

## Abstract

**Ethnopharmacological Relevance:** Gypenosides and *Costus pictus* D.Don are used as an anti-diabetic herbal remedy in China and India respectively. However, the synergistic effect of these two extracts on β-cell protection is not yet elucidated.

**Introduction:** In Type 2 diabetes mellitus (T2DM), pro-inflammatory cytokines and lipotoxicity are known causes of pancreatic β-cell dysfunction and impaired insulin secretion and eventually β-cell death. Thus, any cytoprotective drug supplements can protect the β-cell and may help in T2DM treatment. Gypenosides, extracted from the Chinese medicinal herb *Gynostemma pentaphyllum* and the leaf extract from an Indian medicinal herb *Costus pictus* D. Don are used in traditional medicine due to their insulin secretory properties. In our previous studies, both extracts have shown significant cytoprotective effects in insulin-secreting BRIN-BD11 cells. In the present study, we aim to investigate the synergistic effects of a combination of these extracts on BRIN-BD11 β-cell protection.

**Methods:** Combination of extracts was prepared by adding Gypenosides with *Costus pictus* at 2:1 to a concentration of 18.75mg/ml. Cell viability was determined by MTT assay following treatment with combination and/or palmitate and cytokine cocktail for 24-48h. Following 24h treatment, proliferation was measured by Ki67 staining and cytoprotective gene expression was quantified by qPCR.

**Results:** Combination treatment of 25µg/ml enhanced cell viability both at 24h (n=8; P<0.05) and 48h (n=8; P<0.0001) treatment. Over 24h, combination treatment (25&12.5 µg/ml) showed a significant protective effect against 125µM and 250µM palmitate induced (P<0.0001) and cytokine cocktail-(TNFα 1000U, IL-1β 50U & IFNγ 1000U) (P<0.0001 & P<0.01 respectively) induced toxicity. Combination treatment over 24h increased expression of antioxidant genes Nrf2 (P<0.001), Cat (P<0.001) and Sod1 (P<0.05) along with pro-proliferative Erk1 (P<0.01) while pro-inflammatory Nfkb1 expression was reduced(P<0.001).

**Conclusion:** The results suggest that a combination of gypenosides and *costus pictus* may protect β-cells against inflammatory cytokines and lipotoxicity caused by saturated free fatty acids associated with obesity and diabetes.

## 1. Introduction

Type 2 diabetes mellitus is a complex metabolic disorder identified by insufficient insulin secretion to compensate peripheral insulin resistance or lack of enough insulin-secreting islet β-cells. Although the mechanisms of β-cell dysfunction and loss of β-cells are not clearly understood, high levels of circulating fatty acids, pro-inflammatory cytokines from immune cell infiltration can cause reactive oxygen species accumulation and β-cell demise (Böni-Schnetzler and Meier, 2019; Oh et al., 2018). Thus, any pharmacological agents that could protect β-cell from the detrimental effects of fatty acids and cytokines might have anti-diabetic therapeutic potential. Gypenosides are saponins extracted from the plant *Gynostemma pentaphyllum* and are used as an anti-diabetic herbal tea in china. In our previous studies and various others have reported the insulin secretory properties of gypenosides *in vitro* and *in vivo* (Hoa et al., 2007; Lokman et al., 2015; Patibandla et al., 2015). We also reported the cytoprotective properties of gypenosides in BRIN-BD11 β-cells (Patibandla et al., 2015). *Costus pictus* D.Don also known as the insulin plant is used as an anti-diabetic herbal remedy in India. Leaf extract of *Costus pictus* D.Don is previously reported to have significant insulin secretory properties in cell lines, human islets, and STZ/alloxan-induced animal models (Al-Romaiyan et al., 2010; Gireesh et al., 2009). We have previously reported that *Costus pictus* D.Don secrete both Insulin and GLP-1 in vitro and also have significant β-cell cytoprotective effects against palmitate and cytokine-induced toxicity (Patibandla et al., 2020). In the current study, we have used a combination of gypenosides and *Costus pictus* D.Don leaf extract and elucidated its cytoprotective effect against the detrimental effects of palmitate and cytokine cocktail using clonal BRIN-BD11 β-cell lines.

## 2. Methods

### 2.1 Preparation of Combination

*Costus pictus* D.Don extract was prepared as previously described (Patibandla et al., 2020). Gypenosides (Xi’an Jiatian Biotech Co. Ltd, China, purity 98%) was dissolved in absolute ethanol (25mg/ml) by continuous shaking at room temperature overnight. A combination was prepared by mixing gypenosides and c*ostus pictus* D.Don extract in a 2:1 ratio to get a final concentration of 18.75mg/ml.

### 2.2 Cell culture and viability Assay

BRIN-BD11 cells (a kind gift from Prof. Peter Flatt, Ulster University, Coleraine) were routinely cultured as described previously (Patibandla et al., 2020). Cells were cultured in a 96 well plate (1×10^4^/well) overnight followed by the treatment with combination with or without palmitate or cytokine cocktail. After 24h-48h incubation, cell viability was assessed by MTT assay.

### 2.3 Immunofluorescence

Cells were cultured on glass coverslips (1×10^5^/coverslip) in a 6 well plate overnight followed by 24h treatment with 10nM Exendin-4, 25µg/ml Combination, or vehicle control. The cells were then fixed, permeabilised and stained for Ki67 and insulin as described previously (Patibandla et al., 2020). Primary antibodies used were Anti-Ki67 (AB9260, Merckmillipore, UK) (1:400), Anti-insulin (I-2019, Sigma, UK) (1:200). Secondary antibodies used were goat anti-mouse (Alexa Fluor® 488) (AB150113, abcam, UK) 1:200 and donkey anti-rabbit (Alexa Fluor® 594) (AB150076, Abcam, UK) 1:200. Nuclei were stained with mounting media containing DAPI (Abcam, ab104139). Images were taken using LSM 800 confocal microscope (Carl Zeiss Ltd., Cambridge, UK).

### 2.4 Quantitative real-time PCR

Total RNA was extracted from 24h treated samples using the NucleoSpin® RNA kit (Macherey-Nagel, UK) and cDNA was synthesised using High-Capacity cDNA Reverse Transcription Kit (Applied Biosystems, UK), according to manufacturer’s protocol. Quantitative real-time PCR was performed on a CFX96™ Real-Time PCR detection system using 5X HOT FIREPol® EvaGreen® qPCR Mix Plus (no ROX) (Solis BioDyne, Estonia). Results were analysed by the 2^−ΔΔct^ method.

### 2.5 Statistical Analysis

Results were analysed by Graphpad PRISM® software (Ver. 6.01). Results were presented as mean ± S.E.M. and were analysed by one-way ANOVA with Dunnett’s post-hoc test or Student t-test.

## 3. Results and Discussion

We showed the effects of *costus pictus* D.Don leaf extract and gypenosides on BRIN-BD11 cell viability before and in the current study, the combination of these two increased the % of viable cells over 24h (P<0.05) and over 48h (P<0.0001) (Fig.1A, 1B). Higher concentrations of combination 100-200µg/ml reduced the viability over 24-48h treatment. We further investigated whether this increase in viable cells is due to proliferation by staining with proliferation marker Ki67 and using Exendin-4 (a GLP-1 receptor agonist) as a positive control. As expected, the number of Ki67 positive nuclei is higher in exendin-4 (P<0.05), combination (P<0.05) treated cells (Fig. 2A, 2B). Extracellular signal-regulated kinases (Erk1/2) are previously reported to be associated with β-cell proliferation (Jiang et al., 2018). In parallel to immunofluorescence data, we observed an increase in Erk1 gene expression with combination (P<0.01) treatment (Fig. 2C) Indicating, the combination might promote β-cell proliferation. Proliferative β-cells tend to have reduced Pdx1 and Gck expression (Puri et al., 2018) and our results showed a similar reduction in these gene expressions with treatment (Fig. 2C).

**Figure1:**
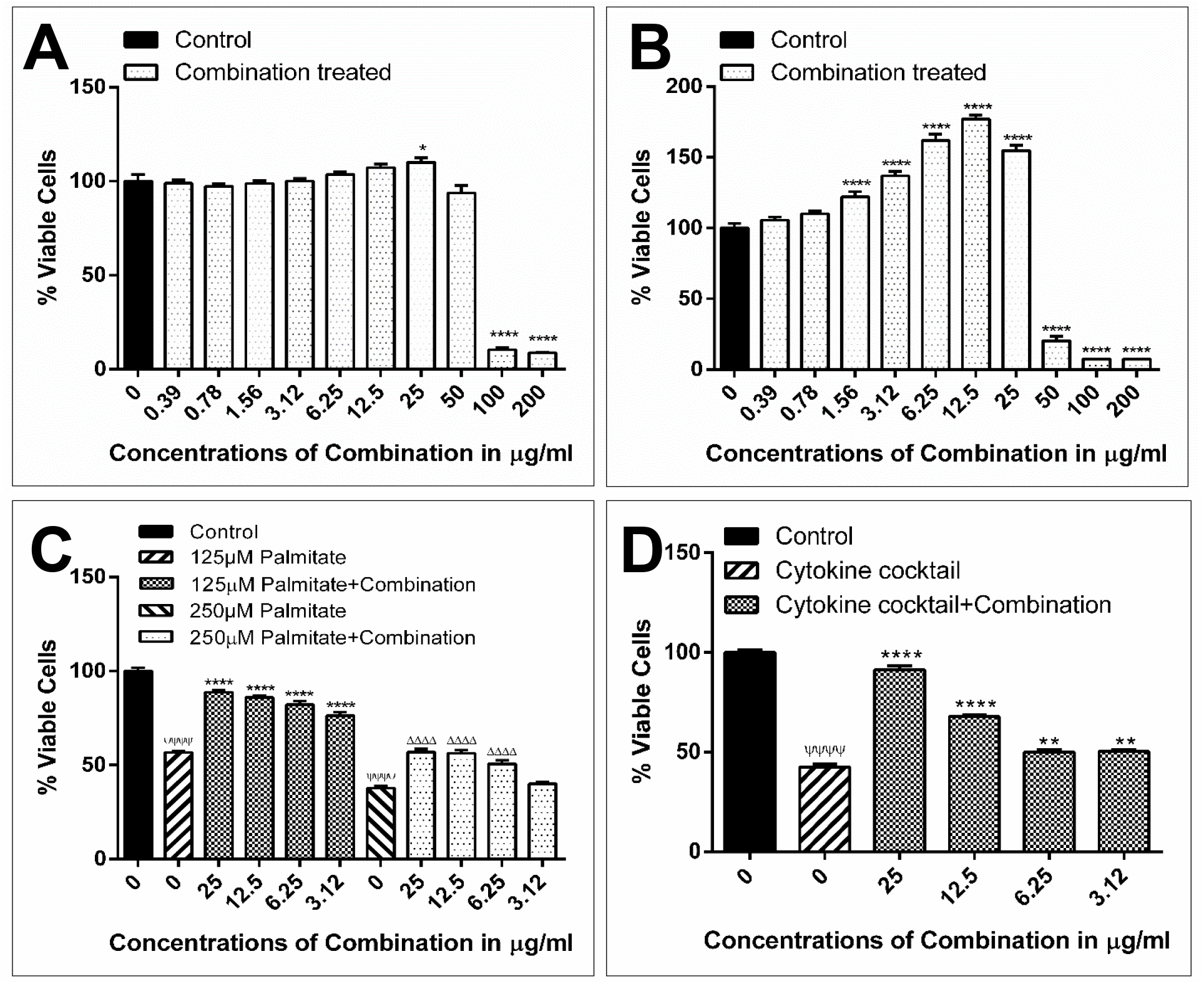
**A&B**, Effects of Combination at concentrations between 200 and 0.39 μg/ml on BRIN-BD11 cell viability over 24 h (**A**) and 48 h (**B**) treatment. **C&D**, Cytoprotective effects of combination against palmitate (**C**) and cytokine cocktail (**D**) induced toxicity. Cells were cultured with palmitate (125/250µM) or cytokine cocktail containing IL-1β (50U), TNF-α (1000U) and IFN-γ (1000U) either alone or along with combination for 24h prior to cell viability measurement by MTT assay. Values represent mean ± S.E.M. from four different experiments conducted in duplicate (n = 4). Data analysed by one-way ANOVA with Dunnett’s posthoc test. *P < 0.05; **P < 0.01; ****P < 0.0001 compared to untreated control (A&B), 125 μM palmitate (C) or cytokine cocktail (D) treatment alone; ΔΔΔΔP<0.0001 compared to 250 μM palmitate(C) alone, ΨΨΨΨP<0.0001, compared to untreated vehicle control (C&D).

**Figure 2:**
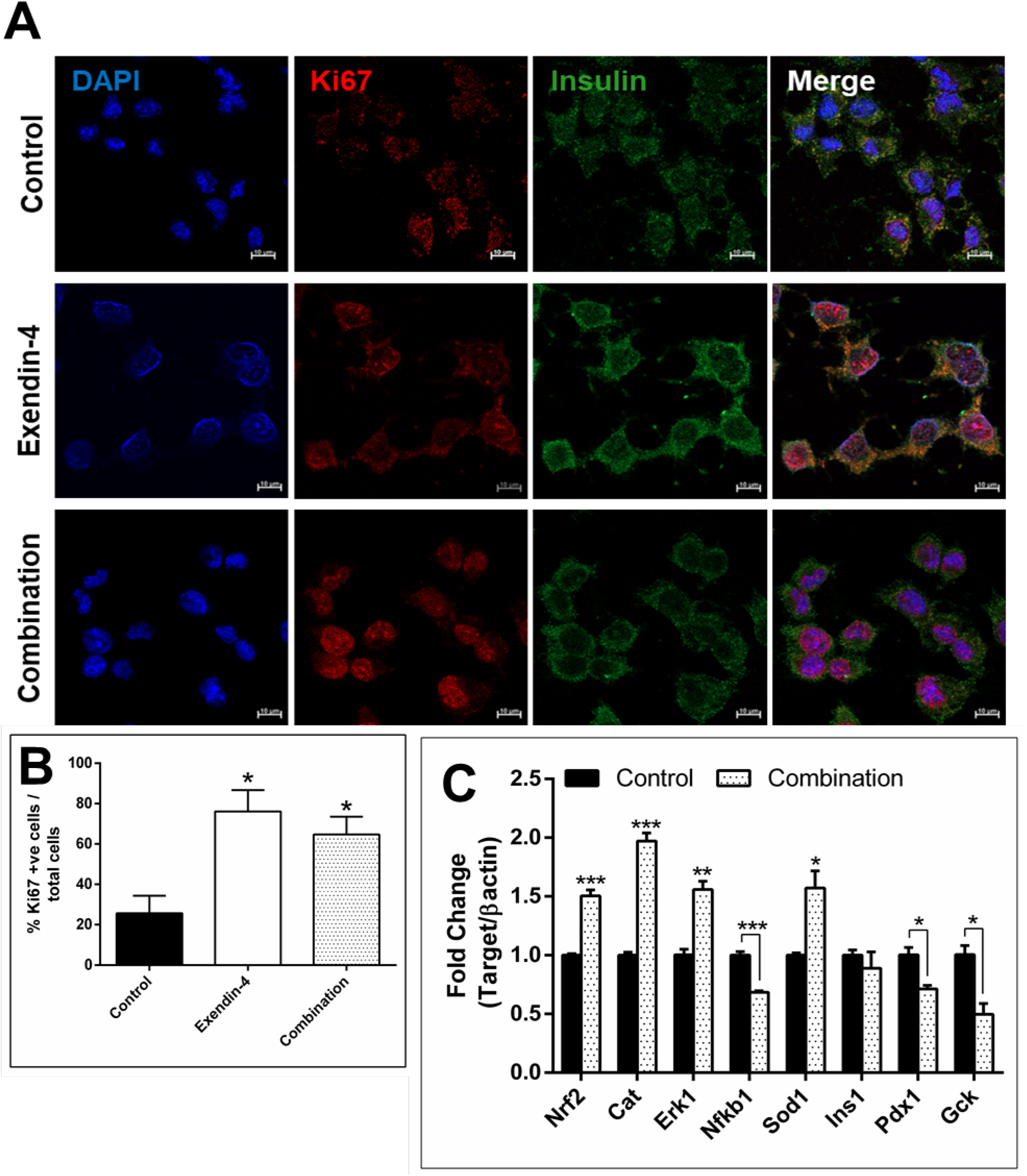
Effects of Combination or Exendin-4 treatment on BRIN-BD11 cell proliferation (**A, B**). BRIN-BD11 cells were treated with 25 μg/ml Combination or Exendin-4 (10nM) for 24h. **A**, Cells were stained with anti-Ki67 (red), Insulin (Green) and nuclear stain DAPI (blue). **B**, Number of Ki67 positive cells were counted from 3 separate experiments and % Ki67 positive cells over total cells was plotted (n = 3). **C**, Effects of Combination (25 μg/ml) treatment over 24h on gene expression. Data represent fold change in mRNA levels compared to vehicle control and normalised to β-actin expression. Values represent mean ± S.E.M. from three different experiments performed in duplicate (n = 3). Data was analysed by one-way ANOVA with Dunnett’s posthoc test (B) or student t-test (C). *P < 0.05, **P < 0.01, ***P < 0.001 compared to their respective control.

High circulating levels of saturated free fatty acids like palmitate are known to cause β-cell dysfunction. Beta cells express G protein-coupled free fatty acid receptors and fatty acid transporter CD36 (Ly et al., 2017). It is known that acutely, activation of fatty acid receptors by palmitate can stimulate insulin secretion. Although, prolonged exposure to palmitate as seen in obesity can cause accumulation of reactive oxygen species and lead to β-cell dysfunction and eventually apoptosis. The combination (6.25-25µg/ml) in the current study significantly (P<0.001) protected the BRIN-BD11 cells against 125 and 250µM palmitate-induced toxicity over 24h treatment (Fig. 1C). Proinflammatory cytokines like IL-1β, TNFα, and IFN-γ are known to cause β-cell inflammation and apoptosis through activating the nuclear factor kappa B (NFκB) pathway (Keane et al., 2015). We used a cocktail of IL-1β (50U), TNF-α (1000U) and IFN-γ (1000U) which reduced the cell viability significantly compared to control (P<0.0001), and addition of the combination 3.12-25µg/ml significantly protected against the cytokine cocktail induced apoptosis (P<0.01-0.0001) (Fig.1D).

Pancreatic β-cells are vulnerable to oxidative stress due to very low levels of antioxidant enzymes expression (Tiedge et al., 1997). Nuclear factor erythroid 2-related factor 2 (Nfe2l2/Nrf2) is the master regulator of antioxidant gene transcription (Abebe et al., 2017). Activation of Nrf2 can regulate gene transcription of superoxide dismutase (Sod1 and Sod2), catalase, and glutathione peroxide 1 (Gpx1) along with many other (Zhu et al., 2005). Intracellular reactive oxygen species are converted into hydrogen peroxide by sod and further to water molecule by catalase. In the current study, we observed a positive correlation of Nrf2(P<0.001), Catalase (P<0.001), and Sod1 (P<0.05) expression in response to 25µg/ml combination (P<0.001) treatment over 24h and Pro-inflammatory Nfkb1 expression was reduced with the combination (P<0.001) treatment (Fig.2C). This might have contributed to its protective effects against palmitate and cytokine cocktail induced cytotoxicity.

## 4. Conclusion

Current results indicate that a combination of gypenosides and costus pictus extract promotes β-cell proliferation and protects against detrimental effects of palmitate and cytokines. Although, further studies are needed to investigate these effects *in vivo*.

## Funding

This research did not receive any specific grant from funding agencies in the public, commercial, or not-for-profit sectors.

## Author contributions

Chinmai Patibandla and Steven Patterson designed experiments. Chinmai Patibandla performed and analysed experiments. Mark James Campbell and Leigh Bennett repeated experiments. Xinhua Shu provided the Gypenoside extract. Chinmai Patibandla and Steven Patterson prepared the manuscript.

## Declarations of interest

none

## Notes

### Competing Interest Statement

The authors have declared no competing interest.

